# Impairments of motor adaptation in Essential Tremor are linked to movement execution

**DOI:** 10.1101/2023.04.21.537795

**Authors:** Flo Blondiaux Pirson, Louisien Lebrun, Bernard J. Hanseeuw, Frédéric Crevecoeur

## Abstract

0.

Essential tremor (ET) is a neurological disorder characterized by involuntary oscillations of the limbs. Previous studies have hypothesized that ET was a cerebellar disorder and reported impairments in motor adaptation. However, recent advances have highlighted that motor adaptation involved several components linked to anticipation and control, all dependent on cerebellum, and the specific alteration of adaptation of ET has not been identified. To address this question we investigated behavioural markers of adaptation in ET patients (n=20) and age-matched healthy volunteers (n=20) in saccadic and upper limb adaptation tasks, probing compensation for target jumps and for velocity-dependent force fields, respectively. We found that both groups adapted their movements to the novel contexts, however, ET patients adapted to a lesser extent compared to healthy volunteers. Importantly, we decomposed movements into components linked to anticipation, preserved here, and real-time execution, which were responsible for the adaptation deficit. Altogether, our results suggest that execution deficits may be a specific functional consequence of the alteration of neural pathways associated with ET.

**Significance Statement:** We tested Essential Tremor patients’ adaptation abilities in classical tasks including saccadic adaptation to target jumps and reaching adaptation to force field disturbances. Patients’ adaptation was present but impaired in both tasks. Interestingly, the deficits were mainly present during movement execution, while the anticipatory components of movements were similar to healthy volunteers. These findings reinforce the hypothesis of a cerebellar origin for essential tremor and details the motor adaptation impairments previously found in this disorder.

## 1. Introduction

Essential tremor (ET) is one of the most common movement disorders provoking involuntary oscillations of patients’ limbs (Louis & Ferreira, 2010). Typical clinical features of ET are kinetic and intention tremors of limbs, particularly upper limbs and, to a lesser extent, head and trunk (Bhatia et al., 2018; Clark & Louis, 2018; Shanker, 2019). To date, the pathophysiology of this neurological disorder is not fully understood, but some previous works have suggested a cerebellar origin (Ibrahim et al., 2021). Neuroimaging studies reported abnormalities in the structure and connectivity of cerebellum as well as differences in the cerebello-thalamo-cortical loop (Mavroudis et al., 2021; Muthuraman et al., 2018; Pietracupa et al., 2021; Tikoo et al., 2020). Postmortem studies also pointed towards alterations of cerebellar cells (Louis & Faust, 2020). However, patients with ET do not suffer from the same intensity of symptoms as patients with cerebellar ataxia, suggesting that some of the main functions of the cerebellum are preserved in this condition.

Previous studies also reported the importance of cerebellum’s integrity in movement control and state estimation (Diedrichsen & Bastian, 2014; McNamee & Wolpert, 2019; Therrien & Bastian, 2015). The central nervous system cannot exactly access the state of the limbs (position, speed, etc.) due to the intrinsic delays in sensory feedback and to sensorimotor noise. As a result of this uncertainty, the brain must predict the next state that will be used to plan and control fast and accurate movements. The prediction is computed based on the delayed sensory feedbacks and an efferent copy of the motor command used in conjunction with prior knowledge of body and environmental dynamics. This operation of state estimation, which makes use of internal models, has been associated with cerebellum (Kilteni et al., 2020; Miall et al., 1993, 2007; Wolpert et al., 1998).

The ability to adapt motor patterns to novel environments rests on our ability to update the internal models used for estimation and control, thereby allows for proactive compensation for sustained disturbance. Accordingly, it has been formulated that adaptation depends on the integrity of cerebellum. The involvement of cerebellum in motor adaptation has been studied with a variety of movements, with non-human species (Baizer et al., 1999; Ojakangas & Ebner, 1992), as well as with cerebellar patients (Smith & Shadmehr, 2005; Tseng et al., 2007; Xu-Wilson et al., 2009), who have shown deficits of adaptation in response to repeated perturbations. These results possibly reflect inaccurate internal models and a relative inability to update them in a way that counters the disturbance induced by the novel environment (Diedrichsen & Bastian, 2014; Therrien & Bastian, 2015).

Given the involvement of cerebellum in sensorimotor adaptation and its putative link with tremor, previous works studied adaptation in cohorts of ET patients. Generally, these patients were able to adapt to perturbations, but to a lesser extent than healthy volunteers in various tasks including adaptation to force field, visuomotor and prism adaptation, as well as eyeblink conditioning (Chen et al., 2006; Bindel et al., 2022; Hanajima et al., 2016; Kronenbuerger et al., 2007). Considering that sensorimotor adaptation may involve multiple components (Mathew & Crevecoeur, 2021), we aimed to better understand the mechanisms at the origin of this motor adaptation deficit using two sensorimotor adaptation tasks probing visual and upper limb motor systems. We studied motor adaptation in saccadic eye movements evoked by peri-saccadic target jumps, and upper limb reaching adaptation to velocity-dependent force fields. We found that motor adaptation deficits observed in both tasks were due to the time course of movement execution while anticipatory compensation for the perturbation was comparable to the control group.

Our results support the hypothesis of a cerebellar origin for essential tremor but suggest a more specific alteration of the pathways linked to the real-time execution of the movement across both visual and upper limb motor systems.

## 2. Methods

### Participants

A total of 22 Essential Tremor (ET) patients and 20 healthy volunteers (HVs) participated in this study. All participants provided written informed consent following procedures approved by the Ethics Committee at the host institution (*Comité d’Éthique Hospitalo Facultaire*, UCLouvain, Belgium). A neurologist performed a clinical evaluation with the Fahn-Tolosa-Marin Tremor Rating Scale (FTM-TRS) (Fahn et al., 1993) on all participants to evaluate the severity of the tremor. ET patients did not interrupt their medication before the assessment. We kept track of current medications and dosages (Supplementary Table 1). All but four ET participants took part in the two experiments. These four patients experienced difficulties with one of the two tasks: inability to perform the task due to their hand tremor, pre-existing shoulder condition, and impossibility to calibrate the eye-tracking system (2 volunteers). As a result, each population of ET patients was composed of 20 volunteers. All HVs participated in the two tasks, resulting in a group of 20 HVs.

The FTM tremor rating scale was divided into three sections. Part A quantified the tremor amplitude of the limbs at rest, with posture holding, and during action. Part B quantified the action tremor of the upper limbs, particularly writing, drawing, and pouring liquids. Part C quantified the functional disability, evaluating patients’ impairments in daily life activities (eating, drinking, working, …). We ignored this part filled by the participant due to the higher subjectivity of some criteria and because our experiments assessed different tasks. Maximum scores for each section were respectively, 80, 32, and 28. Since this scale is not specific to ET evaluation, a low score in some items is expected like rest tremor, uncommon in ET.

### Tasks

The study was composed of two experiments: a saccadic adaptation task and a reaching adaptation task to a force field (Figure 1&3). During the saccadic adaptation task, participants sat in a dark room on a height-adjustable chair in front of a screen. Head movements were restricted using a chin rest bar and forehead support. Eye movements were recorded using an Eyelink 1000 Tower-mounted eye tracker (SR Research, Ottawa, Canada) at a sampling rate of 1kHz, and stored for offline analyses. Participants gazed at the green targets (0.8 visual degrees wide) displayed on the screen. After a series of baseline trials containing a fixation target and a goal target, a jump of the goal target was introduced as a perturbation for the adaptation trials (Figure 1a&b). We divided the experiment into 7 blocks: oblique (30 trials), horizontal (30 trials), and five adaptation blocks (60 trials each).

**Figure 1.**
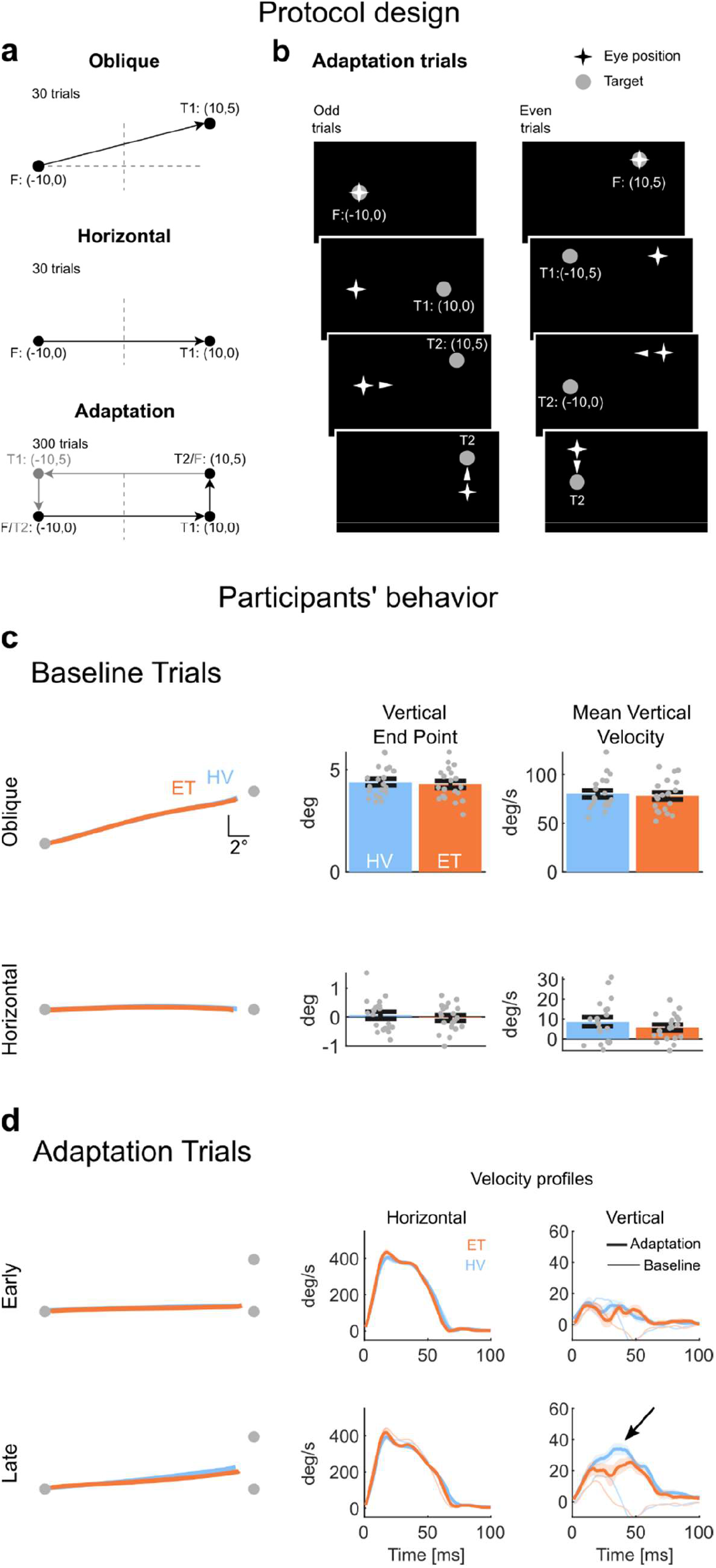
Schematic representation of the saccadic adaptation protocol design (a,b) & behavior in saccadic adaptation task (c,d). **a** Time course of a trial in the saccadic -adaptation task. A: Participants (n=40) were asked to perform saccades to gaze at a target visible on the screen. The experiment was divided into 3 trial types: oblique saccades, horizontal saccades, and adaptation saccades. Each trial began with a fixation target (F), followed by the appearance of the goal target (T1) after a random delay. Locations are indicated in visual degrees. During adaptation trials, the goal target was shifted by 5 degrees (T2). These trials occurred in two directions, to the right (black) and the left (grey). Target jump was counterclockwise. The dashed lines depict the axes centered straight ahead. **b** Target sequence from an adaptation trial. Arrows indicate when a saccade was initiated. Target jump (T1 to T2) occurs when the saccade is initiated. At the end of the trial, T2 served as fixation for the next trial (grey in 1A). **c** (Left) Group average of participants’ eye trajectories averaged over the oblique and horizontal blocks. Grey dots represent the targets. (Right) Vertical end point and mean vertical velocity of saccades, across all trials, and averaged for the two groups. Error bars represent the SEM. Grey dots represent individual datapoints. **d** (Left) Saccades averaged over the first and last 10 trials to highlight differences between early and late adaptation phases. (Right) Average velocity traces are plotted for the horizontal and vertical components. Thin traces correspond to the horizontal baseline trials. Shaded areas represent the SEM.

During the horizontal and oblique trials, a fixation target (F: (−10°,0°)) was projected on the left side of the screen. After a random wait time between 2 sec and 3 sec distributed uniformly, a new target was projected 20° horizontally to the right (Oblique: T:(10°,5°); Horizontal: T: (10°,0°)). The adaptation trials started with the same initial fixation target (F). A second target displayed 20° to the right appeared (T1: (10°,0°)), and participants were instructed to direct their gaze to the new target. As soon as the saccade was detected, the target jumped 5° upward (T2: (10°,5°)). The target T2 served as a fixation target for the next trial, the jump continued to be counterclockwise. The saccades were detected based on a horizontal velocity threshold. Because of the perturbation, the saccade towards T1 was followed by a catch-up saccade towards T2. The analysis focused on the adaptation of the first saccade. An inter trial time of 2.3sec was used in all cases. Participants took a short break between each block (15sec to 1 min).

For the reaching adaptation task, participants sat on a height-adjustable chair in front of an End Point KINARM device (BKIN Technologies, Kingston, Ontario). They grabbed the handle of the robotic arm and performed reaching movements toward the target projected on the screen (Figure 3a). The device recorded the participants’ hand cursor position and forces applied to the handle. An occluder blocked the direct vision of the hand, but a hand-aligned cursor was always represented (white dot, radius 0.6 cm). The radius of the goal target was 1.2 cm. A criterion on movement time was applied to encourage consistent velocities, but all movements crossing the target at least once and finishing within less than 3 cm from its center were kept for analyses. Good movement time ([600 ms – 1500 ms]) was indicated by a green target at the end of the trial. If the participant was too slow, the target remained red and full. For movements too quick, the target switched to red and hollow. The experiment was composed of two trial types: null-field trials, and force field trials. During the null-field block (40 trials), participants were instructed to reach and stop at the displayed target. Two targets were used, equally distant from each other and the fixed start target. The target display order was randomly selected. Each set was composed of the same number of trials towards each target. The adaptation blocks (4 blocks) featured trials with similar instructions but performed in the presence of a clockwise velocity-dependent curl field applied throughout the whole movement. Each block was composed of 66 trials: 30 trials towards each target with the force field, and 3 catch trials per target during which the force field was unpredictably turned off. The trials were presented in random order. The inter-trial time was 1sec. Participants were encouraged to take short breaks between the blocks to reduce fatigue (15 sec to 1 min).

### Data Analysis

Raw measurements of eye position were filtered with a dual-pass 4^th^ order low pass Butterworth filter with a cut-off frequency of 50 Hz. Eye velocity was computed based on position signals using 4^th^ order finite differences. The beginning and end of each saccade were defined using a threshold on the horizontal eye velocity of 16°/s for at least 10 ms. Additional criteria for saccade inclusion were: (1) its duration, within 50 to 200 ms, (2) its peak velocity, above 100 °/s, and (3) a minimum amplitude of 10°, corresponding to one-half of the instructed horizontal displacement. On average, HVs had 11.7% of their trials excluded (min: 3.9%, max: 38.9%), ET patients 14.7% excluded (min: 3.9%, max: 31.4%). Non-detected saccades were mainly due to blinks and fatigue leading to poor detection of the pupil by the eye-tracking system. The mean velocity was computed on a 40ms window, centered around the trial’s peak velocity. The curvature of saccades was approximated using the same method as Xu-Wilson and colleagues (Xu-Wilson et al., 2009). Each saccade was divided into 4 equal segments along the horizontal direction, the slope of the line connecting the start and end of each segment was computed and was referred to as S1-S4 (see Figure 2c-f). After observing no qualitative differences between the left and right saccades, adaptation trials in both directions were combined for analyses.

**Figure 2.**
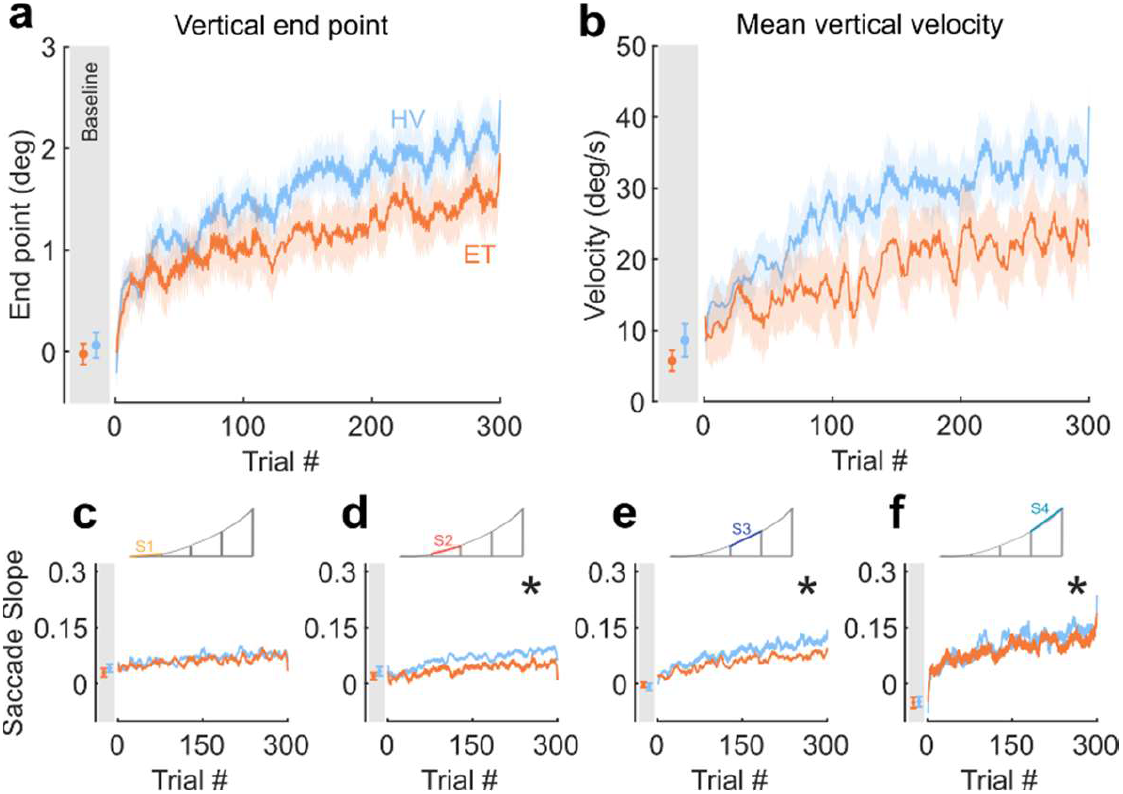
Participants’ behaviour during the saccadic adaptation task. ET patients showed reduced saccadic adaptation **a, b**, Evolution of the vertical end point (a) and peak vertical velocity (b) averaged over each group. Shading indicates SEM. **c-f**, Evolution of saccade curvature with adaptation. The curvature was approximated by dividing saccade into 4 segments and by computing the slope of each segment (labeled S1 – S4). a-f, grey area: Behavior in the horizontal baseline trials, error bars depicts the SEM.

**Figure 3.**
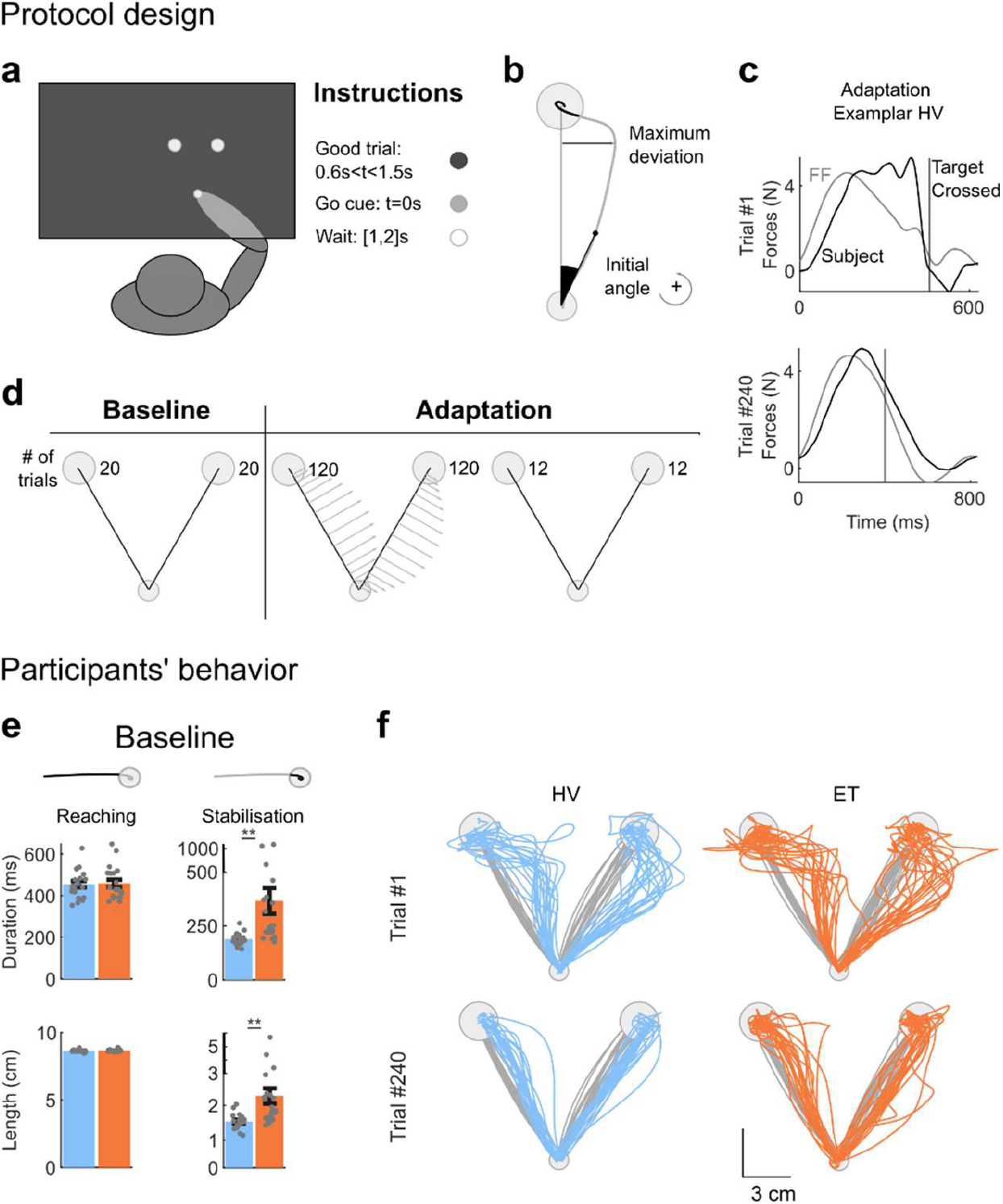
Schematic representation of the upper limb adaptation protocol design (a-d) & Participants behavior during the reaching adaptation task (e,f). **a**, Participants (n=40) were asked to perform reaching movement to one of two goal targets displayed on the screen 10 cm away from the start target with hand-aligned cursor. Direct view of the limb was blocked. The timing of the movement was constrained, the go cue was given by the filling of the goal target. Visual feedback about movement timing was provided (see Methods). **b**, Parameters extracted from individual traces: the initial angle was defined as the angle between the straight line connecting the two targets and the hand position at 200ms. Counter-clockwise angle is defined as positive. Maximum deviation was computed between hand path and the straight line connecting the targets. **c**, Lateral Forces applied by participant on the handle (solid traces) and applied by the robot (grey traces) for an exemplar participant from the control group. Top and bottom panels illustrate the difference between early and late phases of the adaptation blocks. It can be observed that the applied force was closer to the perturbation force at the end of the adaptation phase. **d**, Time course of the protocol: 20 baseline trials towards each target (no force field). Adaptation trials were composed of 120 trials in each direction perturbed by a velocity dependent clockwise curl force field. Randomly interleaved catch trials (no force field) were included during the adaptation blocks. Target presentation order was randomized for each block. **e**, Movements were divided into reaching and stabilization. The end of reaching corresponded to the time at which the cursor crossed the target boundary. Significant differences between the two groups in movement duration and hand path lengths were observed only during the stabilization part. Error bars represent SEM. Note the modified scale for the stabilization measures. **f**, Participants’ hand paths during the first (top) and last (bottom) force field trial. The differences between the trials illustrate adaptation. Grey traces correspond to the last baseline trial of each participant.

For the reaching adaptation task, the data was filtered using a dual-pass 6^th^ order low pass Butterworth filter with a cut-off frequency of 25 Hz. Movement onset and end were defined using a speed threshold of 3cm/s. Movements were included in the dataset if their duration was above 300ms, crossed at least once the target, and ended less than 3cm from the target center. Based on this criterion, we excluded in average 1.1% of the trials for HVs (min: 0.3%, max: 2.3%) and 4.9% of the trials for ET patients (min: 0%, max: 21.1%). Adaptation was quantified using Pearson’s correlation between the force orthogonal to straight path measured at the interface between the hand and the robot handle (measured force), and the force field commanded by the robot and extracted offline based on velocity signals of each trial (commanded force). Intuitively, if the measured force and the commanded force are equal and opposite, one expects high values of correlation between these signals. In contrast, a lack of knowledge of the force field producing only approximate or poorly tuned compensation for the perturbation should yield lower correlation values. It was also verified empirically that this metric was indeed sensitive to adaptation in the same setup and task (Crevecoeur et al., 2020). Initial angles, between the hand path and the straight line connecting the two targets, were computed at 200ms after movement initiation, with a counter-clockwise angle being defined as positive. No qualitative differences were observed between the two targets, movements have therefore been combined for analyses. We used a constant bin width of 8 trials for illustration purposes to show the effect of the perturbations (see Figs 1d – right panel, 2 a, b, e-h, 4 a-e).

### Statistical Analysis

In both experiments, t-tests were used to compare performances between groups during baseline condition (endpoint and maximum velocity of the saccade, movement duration, and path length of the reaching movement) and for the initial angles from catch trials in the reaching adaptation task. T-tests were also used to compare average group age. Adaptation was evaluated using two techniques: exponential fits and Linear Mixed Effects (LME) models. A standard exponential model of adaptation was fitted on group average and 95% confidence intervals of each parameter were compared between ET and HC. The parameter *a* corresponds to the asymptote, *b* is the amplitude of adaptation, and *c* to the learning rate. The variable *i* is the trial number. The exponential fit is as follows:

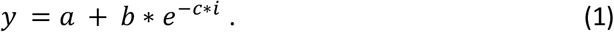

Because the exponential fits were not significant in all cases, we also used linear mixed models to assess the evolution of the dependent variables across trials and group while taking inter-individual variance as a random factor. More precisely, LME models were fitted with factors *Trial Number (i)* and *Group* (2 level factor) as fixed predictors, and a random intercept to capture idiosyncrasy. The significance of each effect was assessed using an F-test and the significance threshold considered was 0.005 (Benjamin et al., 2017). LME and exponential fits were used for vertical end-point, vertical velocity, and saccades slopes in the first experiment, and maximum perturbation forces, normalized deviation, initial angle, and forces correlations in the second experiment. For each dependent variable the formulae of LME was:

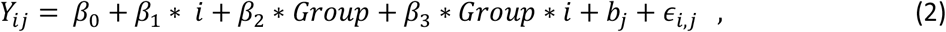

where *Y*_*ij*_ represents the dependent variable of trial number *i* from participant *j, β* are the coefficient of the fixed predictors including trial number, group (control or ET), and interaction between them, and *b*_*j*_ is the random intercept for participant *j*, and *∈*_*i,j*_, is the residual.

Finally, a linear support vector machine classifier was trained on the final performances of each participant in a 2-dimensional space that contained the learning indices that exhibited strong differences across trials and groups. The purpose of the classification technique was to quantify the overlap between the two groups in this two-dimensional space, thus we were interested in the best classifier for the whole dataset and did not perform cross-validation to assess the performance on unseen, test data. In addition to quantifying the overlap between the distribution, the support vector machine was also used to illustrate that the boundary between the two groups separated the plane into overall better or worse performances in the two groups. All analyses were performed offline with Matlab 2020b. The data will be available in DRYARD following the publication of the article.

## 3. Results

### Experiment 1: Saccadic Adaptation

We instructed participants to perform visually guided saccades to a target projected on a screen. The time course of a trial followed standard protocols of saccadic adaptation (Chen-Harris et al., 2008; Xu-Wilson et al., 2009): after an initial fixation delay, the target was projected, and participants initiated a saccade (Figure 1a&b). During movement, the goal target jumped vertically, inducing a terminal error that was gradually reduced as adaptation progressed. A total of 40 volunteers took part to this experiment, 20 Essential Tremor (ET) patients (14 F, 6 M) and 20 Healthy Volunteers (HVs) (14 F, 6 M). There was no significant difference in age between ET patients (M = 58.6, SD = 15.4) and HVs (M=56.8, SD = 12.4); (t-test; t _38_=-0.4013, p= 0.69). We assessed tremor severity in all participants using the Fahn-Tolosa-Marin Tremor Rating Scale (FTM-TRS). ET patients obtained an average score of 8.8 ± 5 (mean ± SD) points in part A assessing tremor amplitude and 14.6 ± 7.6 points for part B which assesses drawings and pouring (part B). The average score of HVs was 1 ± 1.3 (part A) and 1.5 ± 1.7 (part B).

Participants started the experiment with baseline trials, during which the target did not jump. This baseline session was composed of 30 oblique and 30 horizontal saccades, that we later used as reference data for the unperturbed movements. The average eye trajectories are depicted in Figure 1c, left panel. No significant differences of vertical (t-tests; oblique: t_38_=0.47, p=0.64; horizontal: t_38_=0.55, p=0.58) and horizontal (t-tests; oblique: t_38_=0.23, p=0.82; horizontal: t_38_=0.79, p=0.43) end-point were observed between the two participants groups during these baseline trials. Likewise, average vertical (t-tests; oblique: t_38_=0.39, p=0.7; horizontal: t_38_=1.04, p=0.31) and horizontal (t-tests; oblique: t_38_=-0.19, p=0.85; horizontal: t_38_=-0.28, p=0.78) velocities were similar across the two groups. Mean velocity was computed on a 40ms window centered on the trial’s peak velocity. The end-points of patients’ lateral movements were more variable than HVs during oblique trials (Wilcoxon rank-sum test on individual’s variances; Oblique trials: T=312, z=-2.64, p=0.01; Horizontal trials: T=336, z=-1.99, p=0.05). No statistical differences were observed regarding the fixation prior to the saccade, which was assessed by comparing averaged position and velocity across a 100ms window prior saccade initiation.

During the adaptation phase of the experiment, the target jumped vertically (5° up/down) depending on the saccade direction (right/left respectively). The perturbation resulted in an endpoint error corrected by a second saccade. Both groups adapted their saccades to the target jump by increasing the vertical component of the first saccade, but the extent of adaptation was significantly smaller for ET patients. The average eye trajectories at the beginning and the end of adaptation trials are plotted for each participant in Figure 1d. First, the vertical end-point of saccade increased as expected with trial repetitions to reach an average of 2.06° for HVs, and only 1.83° for ET patients as can be seen in Figure 2a. Exponential fits were generally non-significant for the saccadic adaptation task (except for the 3^rd^ slope), we therefore only present LME analyses for this task. A linear mixed-effect (LME) model revealed a significant main effect of the trial (F_(10,356)_ = 24.20, p<10^−4^), no effect of the group (HV or ET) on the vertical end point was observed (F_(38)_ =-0.44, p=0.66) but there was a clear interaction effect between these two factors with a negative coefficient (F_(10,356)_=-5.22, p<10^−4^), showing that ET patients adapted less to the perturbation than healthy volunteers. Likewise, the peak vertical velocity increased with trials for both groups, but with a smaller increase for the ET group (Figure 2b, LME; Trial : F_(10,356)_ = 0.07, P<10^−4^ ; Group: F_(38)_ = -2.16, P= 0.51; Group-by-Trial : F_(10,356)_ = -6.14, P<10^−4^).

Previous studies on saccadic adaptation (Chen-Harris et al., 2008; Xu-Wilson et al., 2009) reported a specific pattern in the evolution of saccades curvature, differentiating adapted saccades from oblique saccades with the same amplitude. We analyzed the evolution of the saccades curvature and observed an altered adaptation pattern in ET patients. We approximated the saccade curvature by four slope segments (S1-S4) (Figure 2c-f, see Methods). The LME model revealed a significant effect of the trial and of the interaction between group and trial for all segments but the first one (LME; S1: Trial: F_(10,356)_ = 8.41, p<10^−4^ ; Group: F_(38)_ = -0.25, p=0.8; Group-by-Trial: F_(10,356)_=-1.25, p=0.21; For S2-S4 Trial: F_(10,356)_ > 15.01, p<10^−4^ ; Group: F_(38)_ > 0.09, p>0.24; Group-by-Trial: F_(10,356)_<-2.88, p<0.004). Differences between HVs and ETs were maximal during S2 and S3. Interestingly, the fact that the first segment of the saccade was similar across groups suggested the preservation of the initial component of saccadic adaptation -which could be seen as a proxy of anticipatory compensation-, whereas differences in S2 and S3 indicated impairment in the time course of saccadic execution supported by internal feedback.

### Experiment 2: Adaptation of reaching movements to a force field

For this experiment, we tested 20 ET patients (15 F, 5 M) and 20 HVs (14 F, 6 M). The HV group was identical to the saccadic adaptation experiment, whereas the ET group involved 18 participants that were also tested in the saccadic adaptation task and 2 new participants. As for the saccadic adaptation task, we verified that there was no significant difference in age between the two groups (ET patients : M: 57.7, SD: 16.1, t-test: t_38_=-0.19, p=0.85). ET patients obtained an FTM-TRS score of 9 ± 4.8 for part A (tremor amplitude) and 13.7 ± 6.5 for part B (drawings/pouring).

Participants first performed 40 movements moving the robotic handle of an instrumented device (KINARM, Kingston, ON) in a null field. These movements were used as control trials to measure differences between the two groups when no perturbation occurred (Figure 3a-d). We decomposed the trials into a reaching and a stabilization phase that were defined based on the time at which participant’s hand crossed the target for the first time. The last movement of each participant in this field is depicted in grey in Figure 3f. The reaching phase was exempt of oscillations: movement duration and path length were similar for both groups (Figure 3e, t-test; Movement duration: t_38_=-0.16, p=0.87; Path length: t_38_=-0.66, p=0.51). The stabilization phase was, however, directly impacted by the tremor with longer movement duration and path length (Figure 3e, t-test; Movement duration: t_38_=-2.95, p=0.005; Path length: t_38_=-3.3, p=0.002). For the subsequent analyses, we focused on the reaching phase of the movement – similar across groups in the absence of perturbation - as adaptation indices taken from this window would likely not be directly affected by the tremor.

When participants were exposed to the force field for the first time, they all experienced large lateral hand deviations (Figure 3f). After a few trials, they quickly adapted their movements to the perturbation, and a reduction of the deviation relative to a straight line was observed: the path length decreased and the forces applied by the participant paralleled the one applied by the robot, a good compensation being the opposite of the perturbation (Figure 3c). Participants exhibited an after-effect when the perturbation was unexpectedly removed, which demonstrated their anticipation and adaptation to the perturbation.

Similar to the saccadic adaptation task, both groups showed an adaptation of their movements to the force field, however, this adaptation was reduced for ET patients. The ET patients were slower in the task, resulting in a significantly smaller perturbation ((Figure 4a); LME: Trial: F_(9255)_ =-13.11, P<10^−4^; Group: F_(38)_ =-1.65, P=0.11 ; Group:Trial: F_(9255)_ =6.62, P<10^−4^). To account for this difference, the maximum deviation measured during each trial (Figure 4b) was normalized by the maximum perturbation force during the trial. The normalized deviation (Figure 4c) was significantly higher in the ET group as revealed by the exponential fits, where all fitted parameters revealed reduced adaptation in the ET population (Figure 4c, asymptote (a) ET: a=0.33 (95% CI: 0.32-0.34), HC: a=0.23 (0.22-0.24), amplitude (b) ET: b=0.30 (0.24-0.37), HC: b=0.21 (0.19-0.23) and learning rate ET: c=0.10 (0.07-0.14), HC: c=0.02 (0.015-0.025)). LME confirmed these results with a significant interaction between the group and the trial number (Figure 4c; LME: Trial: F_(9255)_=-19.45, P<10^−4^; Group: F_(38)_ =0.56, P= 0.58; Group:Trial: F_(9256)_=6.3, P<10^−4^).

**Figure 4.**
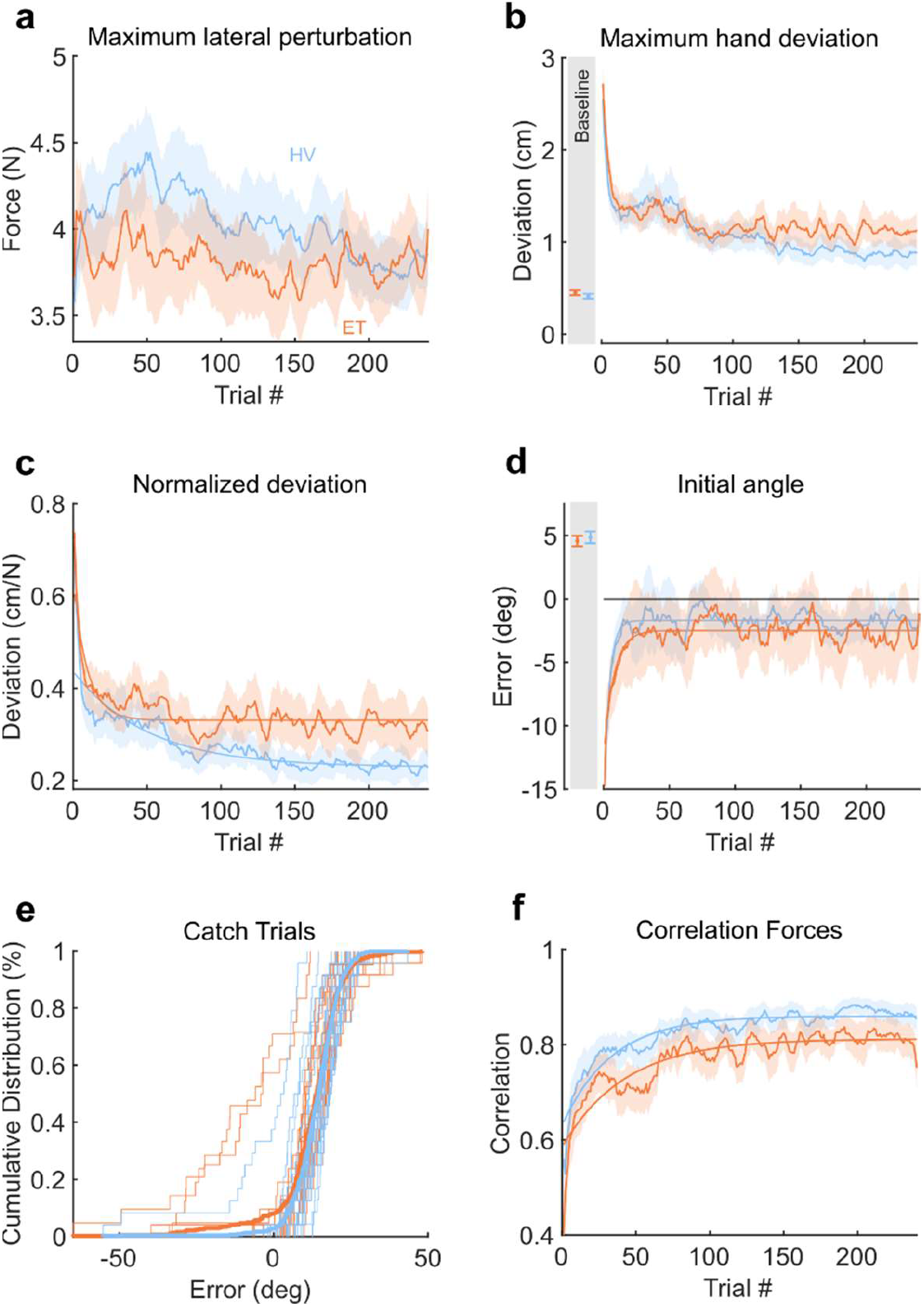
Participants’ adaptation during the reaching task. **a**, Maximum lateral perturbation received by the participants during the adaptation trials. Averaged over each group. **b**, Maximum hand deviation observed during force field trials. **c**, Maximum lateral deviation, normalized by the averaged force received during reaching. **d**, Initial angle between the hand and the line connecting the start-target centers at 200ms after movement initiation. **e**, Cumulative distribution of the initial angle during the catch trials for the ET and healthy participants. **f**, Trial by trial correlation between the force field applied by the robot and the forces applied by the participant. Averaged over participants of each group. **a-d, f**, Shaded areas represent the SEM., grey area: Average behavior in the baseline trials, error bars depicts the SEM. c,d,f: Significant exponential fits.

The initial reach angle, defined as the angle between the position of the hand at 200ms and the straight line connecting the home and the goal target (Figure 3b), can also be studied as a proxy of participants’ anticipation of the perturbation as the earliest modulation of motor commands within movement was previously reported at ∼250ms after movement onset (Crevecoeur et al., 2020; Mathew & Crevecoeur, 2021), suggesting that movement parameters prior to this time must be linked to anticipation. It is expected that the force field produces negative values of initial angles early in the adaptation phase, with gradual changes towards 0 as adaptation progresses. Exponential fits reported similar learning rates and amplitude for both groups but a significant difference in the asymptote (ET: a=-2.5 (95% CI: -2.8,-2.3), HC: a=-1.7 (−1.9,-1.5)). Mixed models analyses reported a small difference in learning rate for the initial angles (Figure 4d), suggesting that both groups had slight differences in initial angles (LME: Trial: F_(9255)_=5.35, P<10^−4^; Group: F_(38)_ =0.03, P=0.98; Group:Trial: F_(9255)_=-3.40, P<10^−3^). However, no difference in initial angles was observed during the catch trials (Figure 4e; t-test; t_38_=0.83, p=0.41), reflecting no significant differences in after-effect between the two groups.

In contrast, the compensation for the force field during movement was more strongly impacted. We computed the correlation between the perturbation force and the force measured at the handle, a high correlation highlighting a good compensation for the force field (Crevecoeur et al., 2020). We found an increase across trials for both groups, while remaining significantly smaller for ET patients than for HVs (Figure 4f). Exponential fits reported a difference of asymptote (ET: a=0.81 (0.8-0.82), HC: a=0.86 (0.85-0.87)) and no significant differences in amplitude or learning rate. This result was also confirmed by LME models (Trial: F_(9021)_=20.44, P<10^−4^; Group: F_(38)_ =-2.41, P=0.021; Trial:Group: F_(9021)_=0.74, P=0.46). Interestingly, the reaching adaptation experiment revealed impairments qualitatively similar to the saccadic adaptation task: first, ET patients were able to adapt, but they exhibited reduced adaptation in comparison with the control group. In both experiments, it appeared that the aspects linked to anticipation were preserved (S1 for the saccadic adaptation, the catch trials initial angles for the reaching task), whereas the metric closely linked to online execution exhibited a reduced compensation for the perturbation (S2 and S3 for the first task, and the correlations computed on continuous force traces for the reaching task).

### Correlations between task performances and FTM-TRS scores

We investigated the correlation of tremor severity with the impairment for each task. Despite ET patients’ significant impairment of saccadic adaptation, the FTM-TRS score (part A+B) was not correlated with the vertical end-point at the end of the adaptation (Figure 5a; Pearson’s correlation = -0.04, p = 0.88). Concerning the force field adaptation task, a relationship between the correlation of forces and the FTM-TRS score was observed (Figure 5b; Pearson’s correlation = -0.52, p = 0.03), suggesting that most affected ET patients were also those showing the larger impairment in reaching adaptation.

**Figure 5.**
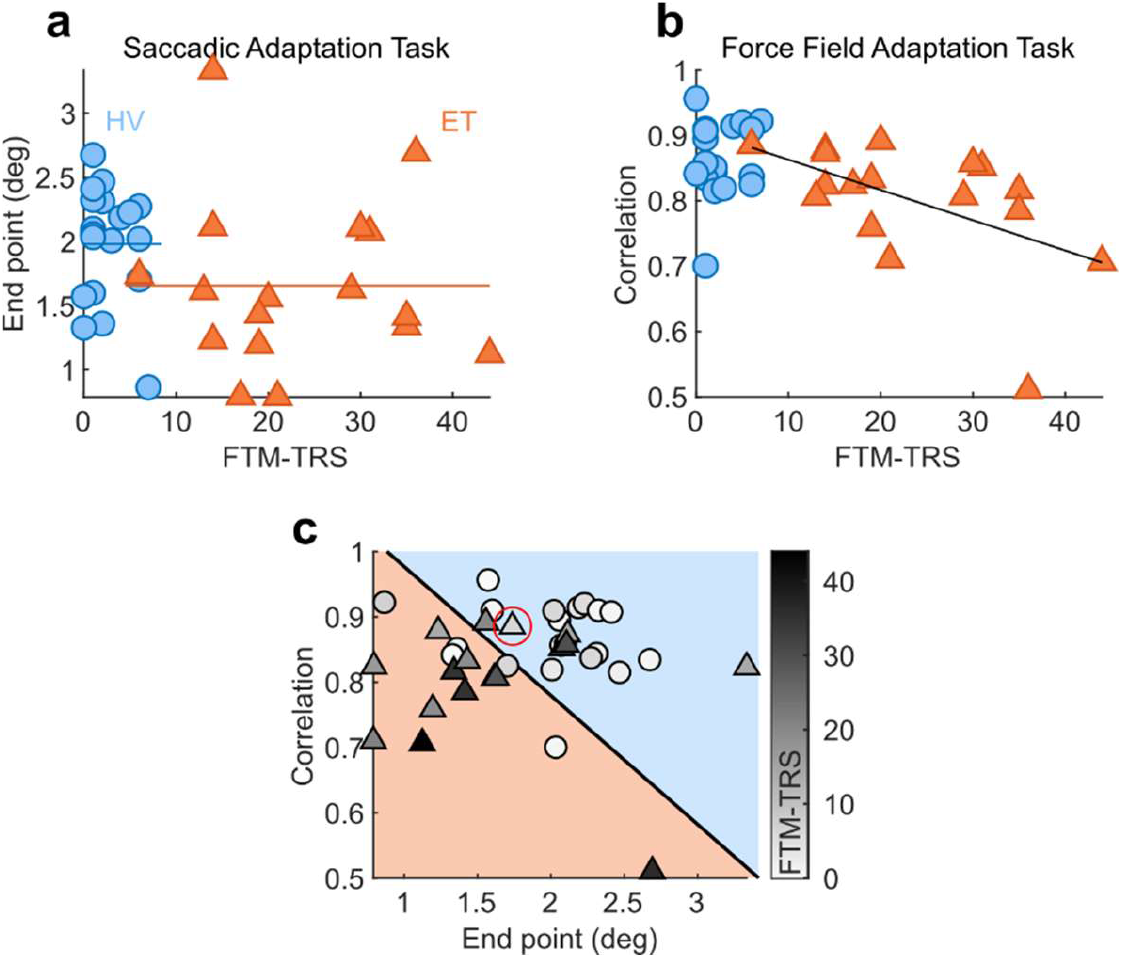
Correlation between performance and tremor score. **a, b**, Relationship between the score obtained during late adaptation (**a**, Saccadic adaptation, **b**, Force field adaptation) and the Tremor Score measured via the FTM TRS scale (Part A+B). Each dot represents the average score obtained over the last 10% trials. **c**, Relationship between the performances in the two tasks for each participant. Color gradient represents the FTM TRS Score. The separation between the two groups was computed by training a linear support vector machine to discriminate populations based on the scores obtained in each task (accuracy: 72.9%). Red circle highlight a misclassification error for a less severely affected ET participant.

Finally, we sought to quantify the differences between the two-dimensional distributions of adaptation indices to measure the overlap across healthy and patient’s groups. We used a linear support vector machine, trained on the vertical end-point error and the continuous correlations, to visualize which task performances regions were associated with each group and quantify the overlap between the two populations taking two-dimensional linear separation in the space of adaptation indices. This method is analogous to Receiver Operating Characteristic in two dimensions. The classifier separated our performance space into reduced performances in both tasks for ET patients and higher performances in both tasks for HVs, as shown in Figure 5c. In this dataset, participants classified in the blue region had an average FTM-TRS score of 7 points; participants classified in the orange region had an average score of 19.5 points. The support vector machine successfully classified 72.9% of the participants. Classification errors mainly occurred for patients with a lower FTM-TRS score. Misclassifications happened more frequently for ET patients with lower FTM-TRS scores, closer to HVs’ scores. An example of misclassification for a less severely affected ET patient is highlighted on Figure 5c.

## 4. Discussion

Our study aimed to better understand the origin of the sensorimotor adaptation deficits previously reported in ET patients. To do so, we selected standard reaching and saccadic adaptation tasks known to be altered in patients with cerebellar deficits. As expected, ET patients experienced a deficit of adaptation in both tasks when compared with the control group. Importantly, our study revealed that the aspects linked to the anticipation of the perturbation, namely the initial saccade slope, the catch trials, and the initial reach angles, clearly evolved across trials and were intact for ET patients. In contrast the deficits seemed to be linked to markers of online control (intermediate saccade slopes and correlations in the reaching task).

Our results are consistent with previous adaptation studies performed with ET patients in various tasks (Bindel et al., 2022; Chen et al., 2006; Hanajima et al., 2016; Kronenbuerger et al., 2007). Here we reproduced the results of Chen et al. (Chen et al., 2006), and added to this line of evidence that the impairment in adaptation in ET patients may reflect a general sensorimotor deficit as it also impacts the visual system despite the absence of ET symptoms in unperturbed saccades (Helmchen et al., 2003; Wójcik-Pȩdziwiatr et al., 2016).

Although a linear correlation was not observed between the tremor score and the performance in the saccadic adaptation task, a correlation was observed between the tremor score and the performance in the force field adaptation task. This result might be explained by the fact that the FTM-TRS scale mostly evaluates upper limb sensorimotor function and tremor, whereas eye movements are not included or evaluated in this clinical scale. The overall tendency was an ordering of the patient’s group in the region of the space of parameters used to quantify adaptation corresponding to lower performances in both tasks.

Our study is not without limitation but they may not have a strong impact on our interpretations. First, participants did not stop their medications before the evaluation, which may influence the FTM-TRS score. Our analyses of the correlations between performance and the tremor score were computed based on this score, and might therefore suffer from the heterogeneity of patients and treatments. Accounting for the medication was however difficult due to the variety of treatments and dosages taken by the participants. Studying sensorimotor deficits while taking medication into account is likely a challenge for prospective studies. Besides the heterogeneity inherent with clinical population and the diversity of treatments, our conclusions are still supported by the fact that the components of motor control prior to the adaptation tasks were similar across groups. Another point of attention concerns the performance of our classifier, which requires validation and should not be hastily generalized. The classifier was used here to quantify the overlap between two distributions, and not to perform prediction on unseen data. The performance of such an approach must be validated with larger datasets and with standard train-test validation before becoming a candidate test to improve the diagnosis of ET. Finally, previous work has reported the existence of cognitive or explicit strategies during adaptation to force fields accounting for a fraction of force produced against a force field (Schween et al., 2020). It is clear that the initial movement angle could be composed of an explicit component, however it was likely similar across groups and our data do not allow us to isolate it. In addition, the main deficit again was linked to movement control which is difficult to relate to a cognitive bias or strategy in both tasks. We believe that quantifying the contribution of explicit components in ET is an interesting topic for follow-up studies.

Cerebellum is suspected to be involved in ET, however, symptoms can vary in nature and intensity from a more generalized disorder like cerebellar ataxia. For instance, patients with cerebellar ataxia showed dramatic impairments and almost complete absence of adaptation in the tasks performed in this present study in comparison with ET patients who showed only reduced adaptation in both tasks (Smith & Shadmehr, 2005; Xu-Wilson et al., 2009). Our motivation to conduct the present study was to parse out specific cerebellar deficits in ET patients. A clear difficulty was that cerebellum has been associated with a wide range of sensorimotor functions, including motor adaptation and control (Baizer et al., 1999; Ojakangas & Ebner, 1992; Smith & Shadmehr, 2005; Tseng et al., 2007; Xu-Wilson et al., 2009). A common ground between adaptation and control is the use of internal models that have been also associated with cerebellum (Diedrichsen & Bastian, 2014; McNamee & Wolpert, 2019; Therrien & Bastian, 2015). It is widely assumed that the brain uses these internal models to produce an estimation of the next state based on the current state estimate and the sensory feedback. This estimation is required to perform fast and accurate movements in an unpredictable environment despite sensorimotor noise and delayed sensory feedback. Online monitoring of the movement and corrections inflight linked to cerebellar mechanisms was shown in saccadic eye movements (Crevecoeur & Kording, 2017; Lefèvre et al., 1998; Quaia et al., 1999; Schreiber et al., 2006; Xu-Wilson et al., 2011), as well as reaching movements (Miall et al., 2007). It is therefore possible that internal models for control hosted in cerebellum would be responsible for tremor and for reduced online compensation in adaptation tasks. Such internal models can take several forms, including aspects linked to sensorimotor coordination, as well as operations linked to timing representations (Bo et al., 2008; Diedrichsen et al., 2007; Kilteni et al., 2019; Martin et al., 1996). In this regard, it is known that oscillations, similar to ET symptoms, can arise in a system that incorrectly compensates for sensorimotor delays (Crevecoeur & Gervers, 2019; Stein & Oĝuztöreli, 1976). Thus, we suggest that tremor arises from incorrect internal models in cerebellum likely associated with delay compensation, producing the well documented oscillations and deficits in online control.

The possible dissociation between anticipation and online control is important for interpreting ET disorders, and also potentially on the underlying dysfunction of the cerebello-thalamo-cortical loop (Helmich et al., 2013; Muthuraman et al., 2018). Our proposed interpretation is that the cerebello-thalamo-cortical pathway conveys an estimate of the state of the system that combines an efferent copy of motor commands with sensory feedback and current estimates that compensates for sensorimotor delays. When the movement was initiated from a static posture, errors had not yet accumulated and movement started in the correct direction, but later in the trial, the faulty delay compensation accumulates errors and the closed-loop system starts producing erroneous control signals potentially leading to oscillations. This view suggests a very specific cause of tremor in ET patients, which is linked to errors in real-time state-estimation based on internal feedback of motor commands. Orienting future research on online feedback control in ET should bring novel insights to validate or invalidate the hypothesis of impaired delay compensation as candidate root cause of oscillations in this population.

To conclude, our contribution is twofold: on the one hand we replicate the presence of cerebellar-dependent adaptation deficits in ET population, adding to the line of evidence that this disorder is linked to cerebellar dysfunction. On the other hand, we proposed a movement decomposition suggesting a specific functional consequence of the altered cerebellar pathways, which affects online control more than anticipation.

## Acknowledgments

Florence Blondiaux is a FRIA grantee of the Fonds de la Recherche Scientifique – FNRS (Be). The FNRS provided salary and research support for Bernard Hanseeuw under grants n° CCL40010417 and n° FRFS-WELBIO40010035. Frédéric Crevecoeur is supported by a grant from the FNRS under grant number 1.C.033.18.

## 6. Supplementary material

**Supplementary Table 1:** Medication usually prescribed to reduce symptoms in ET.

|  | ET | HV |
| --- | --- | --- |
| Propranolol | 9 | 0 |
| Beta Blockers (Other than Propranolol) | 2 | 3 |
| Primidone | 2 | 0 |
| Propranolol + Primidone | 3 | 0 |
| No Beta Blockers nor Primidone | 6 | 17 |
| **Number of participants** | **22** | **20** |

## References

Baizer, J. S., Kralj-Hans, I., & Glickstein, M. (1999). Cerebellar Lesions and Prism Adaptation in Macaque Monkeys. Journal of Neurophysiology, 81(4), 1960–1965. https://doi.org/10.1152/jn.1999.81.4.1960

Benjamin, D. J., Berger, J. O., Johannesson, M., Nosek, B. A., Wagenmakers, E.-J., Berk, R., Bollen, K. A., Brembs, B., Brown, L., Camerer, C., Cesarini, D., Chambers, C. D., Clyde, M., Cook, T. D., De Boeck, P., Dienes, Z., Dreber, A., Easwaran, K., Efferson, C., … Johnson, V. E. (2017). Redefine statistical significance. Nature Human Behaviour, 2(1), 6–10. https://doi.org/10.1038/s41562-017-0189-z

Bhatia, K. P., Bain, P., Bajaj, N., Elble, R. J., Hallett, M., Louis, E. D., Raethjen, J., Stamelou, M., Testa, C. M., Deuschl, G., & the Tremor Task Force of the International Parkinson and Movement Disorder Society. (2018). Consensus Statement on the classification of tremors. from the task force on tremor of the International Parkinson and Movement Disorder Society: IPMDS Task Force on Tremor Consensus Statement. Movement Disorders, 33(1), 75–87. https://doi.org/10.1002/mds.27121

Bindel, L., Mühlberg, C., Pfeiffer, V., Nitschke, M., Müller, A., Wegscheider, M., Rumpf, J.-J., Zeuner, K. E., Becktepe, J. S., Welzel, J., Güthe, M., Classen, J., & Tzvi, E. (2022). Visuomotor Adaptation Deficits in Patients with Essential Tremor. The Cerebellum. https://doi.org/10.1007/s12311-022-01474-5

Bo, J., Block, H. J., Clark, J. E., & Bastian, A. J. (2008). A Cerebellar Deficit in Sensorimotor Prediction Explains Movement Timing Variability. Journal of Neurophysiology, 100(5), 2825–2832. https://doi.org/10.1152/jn.90221.2008

Chen, H., Hua, S. E., Smith, M. A., Lenz, F. A., & Shadmehr, R. (2006). Effects of human cerebellar thalamus disruption on adaptive control of reaching. Cerebral Cortex, 16(10), 1462–1473. https://doi.org/10.1093/cercor/bhj087

Chen-Harris, H., Joiner, W. M., Ethier, V., Zee, D. S., & Shadmehr, R. (2008). Adaptive control of saccades via internal feedback. Journal of Neuroscience, 28(11), 2804–2813. https://doi.org/10.1523/JNEUROSCI.5300-07.2008

Clark, L. N., & Louis, E. D. (2018). Essential tremor. In Handbook of Clinical Neurology (Vol. 147). Elsevier B.V. https://doi.org/10.1016/B978-0-444-63233-3.00015-4

Crevecoeur, F., & Gervers, M. (2019). Filtering compensation for delays and prediction errors during sensorimotor control. 2954, 2925–2954. https://doi.org/10.1162/NECO

Crevecoeur, F., & Kording, K. P. (2017). Saccadic suppression as a perceptual consequence of efficient sensorimotor estimation. ELife, 6, 1–15. https://doi.org/10.7554/eLife.25073

Crevecoeur, F., Thonnard, J. L., & Lefèvre, P. (2020). A very fast time scale of human motor adaptation: Within movement adjustments of internal representations during reaching. ENeuro, 7(1), 1–16. https://doi.org/10.1523/ENEURO.0394-19.2019

Diedrichsen, J., & Bastian, A. J. (2014). Cerebellar Function. The Cognitive Neurosciences, 451.

Diedrichsen, J., Criscimagna-Hemminger, S. E., & Shadmehr, R. (2007). Dissociating Timing and Coordination as Functions of the Cerebellum. Journal of Neuroscience, 27(23), 6291–6301. https://doi.org/10.1523/JNEUROSCI.0061-07.2007

Fahn, S., Tolosa, E., & Marin, C. (1993). Clinical Rating Scale for Tremor. Parkinson’s Disease and Movement Disorders, 2, 271–280.

Hanajima, R., Tsutsumi, R., Shirota, Y., Shimizu, T., Tanaka, N., & Ugawa, Y. (2016). Cerebellar dysfunction in essential tremor. Movement Disorders, 31(8), 1230–1234. https://doi.org/10.1002/mds.26629

Helmchen, C., Hagenow, A., Miesner, J., Sprenger, A., Rambold, H., Wenzelburger, R., Heide, W., & Deuschl, G. (2003). Eye movement abnormalities in essential tremor may indicate cerebellar dysfunction. Brain, 126(6), 1319–1332. https://doi.org/10.1093/brain/awg132

Helmich, R. C., Toni, I., Deuschl, G., & Bloem, B. R. (2013). The pathophysiology of essential tremor and parkinson’s tremor. Current Neurology and Neuroscience Reports, 13(9). https://doi.org/10.1007/s11910-013-0378-8

Ibrahim, M. F., Beevis, J. C., & Empson, R. M. (2021). Essential Tremor –A Cerebellar Driven Disorder? Neuroscience, 462(November), 262–273. https://doi.org/10.1016/j.neuroscience.2020.11.002

Kilteni, K., Engeler, P., & Ehrsson, H. H. (2020). Efference Copy Is Necessary for the Attenuation of Self-Generated Touch. IScience, 23(2), 100843. https://doi.org/10.1016/j.isci.2020.100843

Kilteni, K., Houborg, C., & Ehrsson, H. H. (2019). Rapid learning and unlearning of predicted sensory delays in self-generated touch. ELife, 8, e42888. https://doi.org/10.7554/eLife.42888

Kronenbuerger, M., Gerwig, M., Brol, B., Block, F., & Timmann, D. (2007). Eyeblink conditioning is impaired in subjects with essential tremor. Brain, 130(6), 1538–1551. https://doi.org/10.1093/brain/awm081

Lefèvre, P., Quaia, C., & Optican, L. M. (1998). Distributed model of control of saccades by superior colliculus and cerebellum. Neural Networks, 11(7–8), 1175–1190. https://doi.org/10.1016/S0893-6080(98)00071-9

Louis, E. D., & Faust, P. L. (2020). Essential tremor pathology: Neurodegeneration and reorganization of neuronal connections. Nature Reviews Neurology, 16(2), 69–83. https://doi.org/10.1038/s41582-019-0302-1

Louis, E. D., & Ferreira, J. J. (2010). How common is the most common adult movement disorder? Update on the worldwide prevalence of essential tremor. Movement Disorders, 25(5), 534–541. https://doi.org/10.1002/mds.22838

Martin, T. A., Keating, J. G., Goodkin, H. P., Bastian, A. J., & Thach, W. T. (1996). Throwing while looking through prisms: I. Focal olivocerebellar lesions impair adaptation. Brain, 119(4), 1183–1198. https://doi.org/10.1093/brain/119.4.1183

Mathew, J., & Crevecoeur, F. (2021). Adaptive Feedback Control in Human Reaching Adaptation to Force Fields. Frontiers in Human Neuroscience, 15, 742608. https://doi.org/10.3389/fnhum.2021.742608

Mavroudis, I., Petrides, F., Karantali, E., Chatzikonstantinou, S., McKenna, J., Ciobica, A., Iordache, A. C., Dobrin, R., Trus, C., & Kazis, D. (2021). A voxel-wise meta-analysis on the cerebellum in essential tremor. Medicina (Lithuania), 57(3), 1–10. https://doi.org/10.3390/medicina57030264

McNamee, D., & Wolpert, D. M. (2019). Internal Models in Biological Control. Annual Review of Control, Robotics, and Autonomous Systems, 2(1), 339–364. https://doi.org/10.1146/annurev-control-060117-105206

Miall, R. C., Christensen, L. O. D., Cain, O., & Stanley, J. (2007). Disruption of state estimation in the human lateral cerebellum. PLoS Biology, 5(11), 2733–2744. https://doi.org/10.1371/journal.pbio.0050316

Miall, R. C., Weir, D. J., Wolpert, D. M., & Stein, J. F. (1993). Is the Cerebellum a Smith Predictor? Journal of Motor Behavior, 25(3), 203–216. https://doi.org/10.1080/00222895.1993.9942050

Muthuraman, M., Raethjen, J., Koirala, N., Anwar, A. R., Mideksa, K. G., Elble, R., Groppa, S., & Deuschl, G. (2018). Cerebello-cortical network fingerprints differ between essential, Parkinson’s and mimicked tremors. Brain, 141(6), 1770–1781. https://doi.org/10.1093/brain/awy098

Ojakangas, C. L., & Ebner, T. J. (1992). Purkinje cell complex and simple spike changes during a voluntary arm movement learning task in the monkey. Journal of Neurophysiology, 68(6), 2222–2236. https://doi.org/10.1152/jn.1992.68.6.2222

Pietracupa, S., Bologna, M., Tommasin, S., Berardelli, A., & Pantano, P. (2021). The Contribution of Neuroimaging to the Understanding of Essential Tremor Pathophysiology: A Systematic Review. Cerebellum, 0123456789. https://doi.org/10.1007/s12311-021-01335-7

Quaia, C., Lefèvre, P., & Optican, L. M. (1999). Model of the Control of Saccades by Superior Colliculus and Cerebellum. Journal of Neurophysiology, 82(2), 999–1018. https://doi.org/10.1152/jn.1999.82.2.999

Schreiber, C., Missal, M., & Lefèvre, P. (2006). Asynchrony Between Position and Motion Signals in the Saccadic System. Journal of Neurophysiology, 95(2), 960–969. https://doi.org/10.1152/jn.00315.2005

Schween, R., McDougle, S. D., Hegele, M., & Taylor, J. A. (2020). Assessing explicit strategies in force field adaptation. Journal of Neurophysiology, 123(4), 1552–1565. https://doi.org/10.1152/jn.00427.2019

Shanker, V. (2019). Essential tremor: Diagnosis and management. The BMJ, 366. https://doi.org/10.1136/bmj.l4485

Smith, M. A., & Shadmehr, R. (2005). Intact ability to learn internal models of arm dynamics in Huntington’s disease but not cerebellar degeneration. Journal of Neurophysiology, 93(5), 2809–2821. https://doi.org/10.1152/jn.00943.2004

Stein, R. B., & Oguztöreli, M. N. (1976). Tremor and other oscillations in neuromuscular systems. Biological Cybernetics, 22(3), 147–157. https://doi.org/10.1007/BF00365525

Therrien, A. S., & Bastian, A. J. (2015). Cerebellar damage impairs internal predictions for sensory and motor function. Current Opinion in Neurobiology, 33, 127–133. https://doi.org/10.1016/j.conb.2015.03.013

Tikoo, S., Pietracupa, S., Tommasin, S., Bologna, M., Petsas, N., Bharti, K., Berardelli, A., & Pantano, P. (2020). Functional disconnection of the dentate nucleus in essential tremor. Journal of Neurology, 0123456789. https://doi.org/10.1007/s00415-020-09711-9

Tseng, Y. W., Diedrichsen, J., Krakauer, J. W., Shadmehr, R., & Bastian, A. J. (2007). Sensory prediction errors drive cerebellum-dependent adaptation of reaching. Journal of Neurophysiology, 98(1), 54–62. https://doi.org/10.1152/jn.00266.2007

Wójcik-Pędziwiatr, M., Plinta, K., Krzak-Kubica, A., Zajdel, K., Falkiewicz, M., Dylak, J., Ober, J., Szczudlik, A., & Rudzińska, M. (2016). Eye movement abnormalities in essential tremor. Journal of Human Kinetics, 52(1), 53–64. https://doi.org/10.1515/hukin-2015-0193

Wolpert, D. M., Miall, R. C., & Kawato, M. (1998). Internal models in the cerebellum. Trends in Cognitive Sciences, 2(9), 338–347. https://doi.org/10.1016/S1364-6613(98)01221-2

Xu-Wilson, M., Chen-Harris, H., Zee, D. S., & Shadmehr, R. (2009). Cerebellar contributions to adaptive control of saccades in humans. Journal of Neuroscience, 29(41), 12930–12939. https://doi.org/10.1523/JNEUROSCI.3115-09.2009

Xu-Wilson, M., Tian, J., Shadmehr, R., & Zee, D. S. (2011). TMS Perturbs Saccade Trajectories and Unmasks an Internal Feedback Controller for Saccades. Journal of Neuroscience, 31(32), 11537–11546. https://doi.org/10.1523/JNEUROSCI.1584-11.2011

